# Non-canonical D1-D2 recombination produces two types of rearrangements and represents a conservative hidden stage of T-cell receptor beta chain generation

**DOI:** 10.1101/2023.04.21.537904

**Authors:** Anastasia O. Smirnova, Anna M. Miroshnichenkova, Laima D. Belyaeva, Ilya V. Kelmanson, Yuri B. Lebedev, Ilgar Z. Mamedov, Dmitriy M. Chudakov, Alexander Y. Komkov

**Affiliations:** Central European Institute of Technology, Masaryk University, Brno, Czech Republic; Shemyakin-Ovchinnikov Institute of Bioorganic Chemistry, Moscow, Russia; Abu Dhabi Stem Cells Center (ADSCC), Abu Dhabi, United Arab Emirates; Pirogov Russian National Research Medical University, Moscow, Russia; Weizmann Institute of Science, Rehovot, Israel; Dmitry Rogachev National Medical and Research Center of Pediatric Hematology, Oncology, and Immunology, Moscow, Russia

## Abstract

T-cell receptor (TCR) diversity is generated by VDJ recombination. The classical course of TCR beta (TRB) chain production starts with D and J segment recombination and finishes with subsequent recombination between the resulting DJ junction and V segment. In this study, we performed deep sequencing of poorly explored incomplete TRBD1 to TRBD2 rearrangements in T-cell genomic DNA. We reconstructed full repertoires of human incomplete TRB DD rearrangements and validated its authenticity by detecting excision circles with RSS (recombination signal sequence) junctions for the first time. The identified rearrangements generated in compliance with the classical 12/23 rule are common for humans, rats, and mice and contain typical VDJ recombination footprints. Detected bimodal distribution of DD junctions indicates two active recombination sites producing long and short DD rearrangements. Unlike long DD rearrangements, the short ones have unusual origin resulting from non-canonical intrachromosomal RSSs’ junctions formation. Identified DD rearrangements lead to deleting J1 and C1 segments and creating diverse hybrid D segments, which recombine further with J2 and V segments. Resulting functional TRB VDDJ rearrangements are present in the memory T-cells subset proving its participation in antigen recognition.

## Introduction

T-cell receptor beta chains (TRB) are one of the essential components of the antigen recognition complex in *αβ*T cells. The power of TCR to recognize a wide variety of possible exterior pathogen peptides is provided by the enormous diversity of both alpha and beta chains. This diversity is a result of a somatic process called VDJ recombination in which the particular V (variable), D (diversity), and J (joining) segments from cassettes located in the TRB locus join together to form mature TRB rearrangement. The recombination process is accompanied by deleting and inserting a random number of random nucleotides in the segment’s junctions. Semi-random choice of segments from the cassettes and random junctions’ formation are the main sources of the required TRB diversity. The classical model of the VDJ recombination for TRB locus postulates the two sequential stages of rearrangement: D to J joining and subsequent V to DJ joining (Schatz and Ji, 2011). The final part of TRB is the C segment which joins VDJ rearrangement at the RNA level via splicing. According to the IMGT database (Giudicelli et al., 2005) human TRB locus contains 48 functional V segments, two D segments, 13 J segments, and two C segments. Importantly, D, J, and C segments are organized into two distinct groups. The first group contains D1, J1.1-1.7, and C1 segments; the second group contains D2, J2.1-2.6, and C2 segments (Figure 1B). The first group is located between V segments and the second group. Each V, D, and J segment are flanked by recombination signal sequences (RSS).

**Figure 1.**
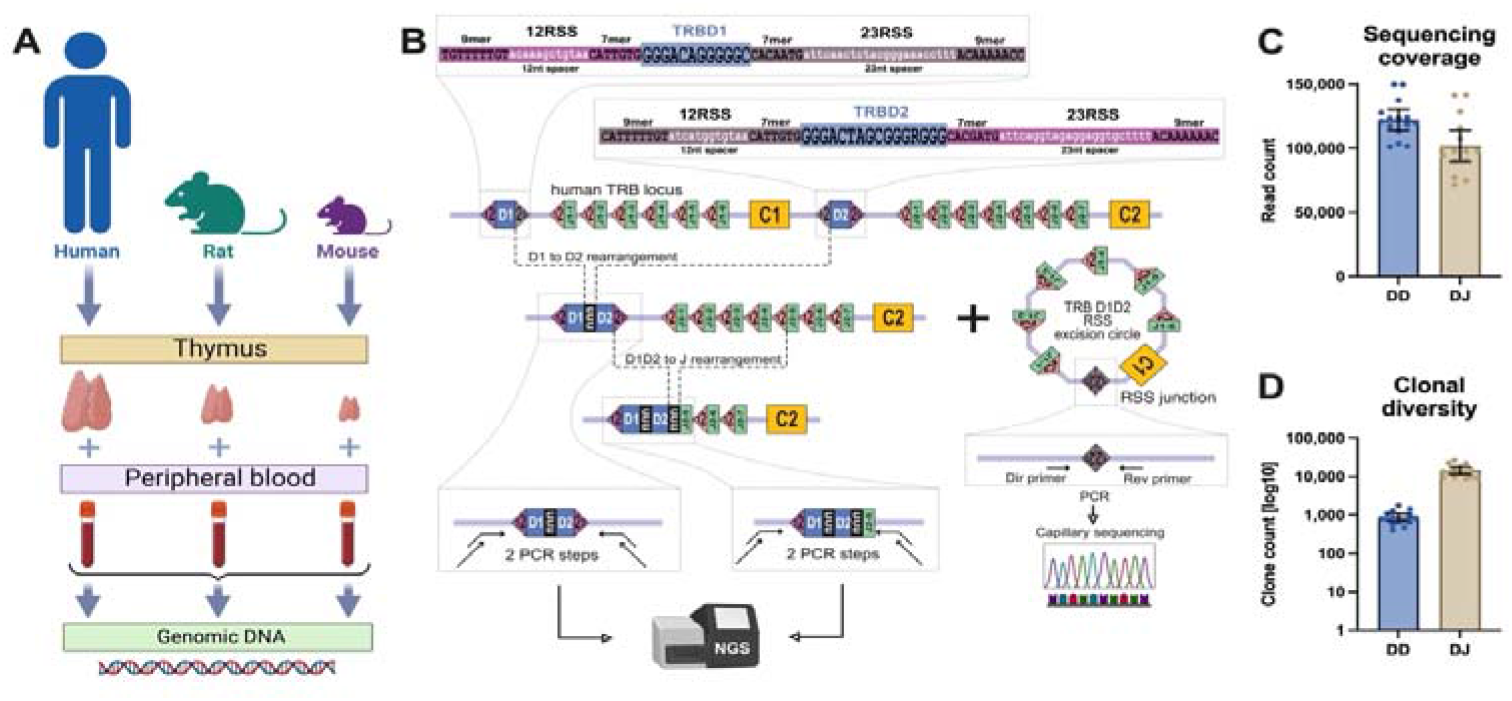
TRB DD and DJ rearrangement sequencing. **A**. Type of collected samples. **B**. PCR-based library preparation and RSS excision circle detection schemes. **C**. Obtained sequencing coverage for TRB DD and TRB DJ libraries. **D**. Clonal diversity of detected TRB DD and DJ rearrangements per ∼100,000 PBMCs. Bar and whiskers represent the mean with a 95% confidence interval.

There are two canonical types of RSS: 12RSS (contains conservative 7-mer and 9-mer separated by less conservative 12 nucleotide spacer) and 23RSS (contains conservative 7-mer and 9-mer separated by less conservative 23 nucleotide spacer). The recombination is initiated by binding RAG-1 and RAG-2 recombinases to RSS signals, leading to segment synapsis formation with subsequent specific DNA cleavage and hairpin generation at the segment’s edges (Schatz and Swanson, 2011). Another recruited protein, Artemis endonuclease, opens DNA hairpins by single strand cut in a random position near segment ends. Exonuclease removes, and at the same time terminal deoxynucleotidyl transferase (TdT) adds several random nucleotides to single-strand ends produced by Artemis. At the final stage, polymerase fills the gaps, and ligase IV repairs the breaks. Because of the binding manner of the RAG-1/RAG-2 complex (Ciubotaru et al., 2015), the VDJ recombination can proceed solely between different RSS types but not between the same ones (12/23 rule) (Eastman et al., 1996). V segments are one-side flanked by 23RSS, J segments are similarly one-side flanked by 12RSS, and D segments are two-side flanked by both 23RSS and 12RSS. Therefore the 12/23 rule does not prohibit direct recombination between D1 and D2 segments. Moreover, sporadic TRB DD rearrangements have indeed been detected in mice previously (Curry and Schlissel, 2008; Hempel et al., 1998). However, for humans, only the TCR delta locus has been analyzed in more or less detail regarding DD rearrangements (Biondi et al., 1990). Here using the advances of high-throughput sequencing, we obtained and comprehensively characterized full repertoires of TRB DD rearrangements in human, mouse, and rat genomes for the first time.

## Results

### DD rearrangement detection at the DNA level

The previous studies (Liu et al., 2014; Safonova and Pevzner, 2020) demonstrated that the conventional concept of VDJ recombination should be reevaluated regarding the detected fraction of non-canonical VDDJ rearrangements. In this study, we performed deep sequencing and analysis of the possible source of VDDJ rearrangements in TRB locus – incomplete DD rearrangements. The analysis was performed at the level of DNA extracted from human, rat, and mouse thymes and blood cells (Figure 1A). Using high-throughput sequencing of amplicons obtained in PCR with primers annealing upstream of 12RSS of the TRB D1 segment and downstream of 23RSS of the TRB D2 segment (Figure 1B), we successfully identified repertoires of incomplete TRB D1-D2 genomic rearrangements.

These DD rearrangements were detected in all expected DNA samples from thymic cells and peripheral blood mononuclear cells (PBMC) of all three analyzed species (Figure 1A). These types of rearrangements were absent in negative control DNA from human CD19+ B cells and non-lymphoid cell line RMS.

In parallel with DD rearrangements, we additionally identified well-known incomplete TRB DJ rearrangements for the same human DNA samples. Despite the similar sequencing coverages (Figure 1C), the clonal diversity of TRB DD rearrangements was ten times less than that of DJ rearrangements (Figure 1D). However, the absolute number of different identified DD clonotypes (1,000 per 100,000 cells average) shows that these rearrangements have been subjected to diversification process during the formation. Typical detected DD rearrangement contained 12RSS of the TRB D1 segment and 23RSS of the TRB D2 segment connected by extremely diverse nucleotide sequences only partially matched to the genome (Figure 2A). These various junctions contain random non-genomic nucleotide sequences flanked by fragments of TRB D1 and D2 segments in nearly half of the cases of detected DD rearrangements. The presence of random non-templated nucleotides and cut D1 and D2 segments are the main source of clonal diversity and, simultaneously, are the hallmark of the VDJ recombination process.

**Figure 2.**
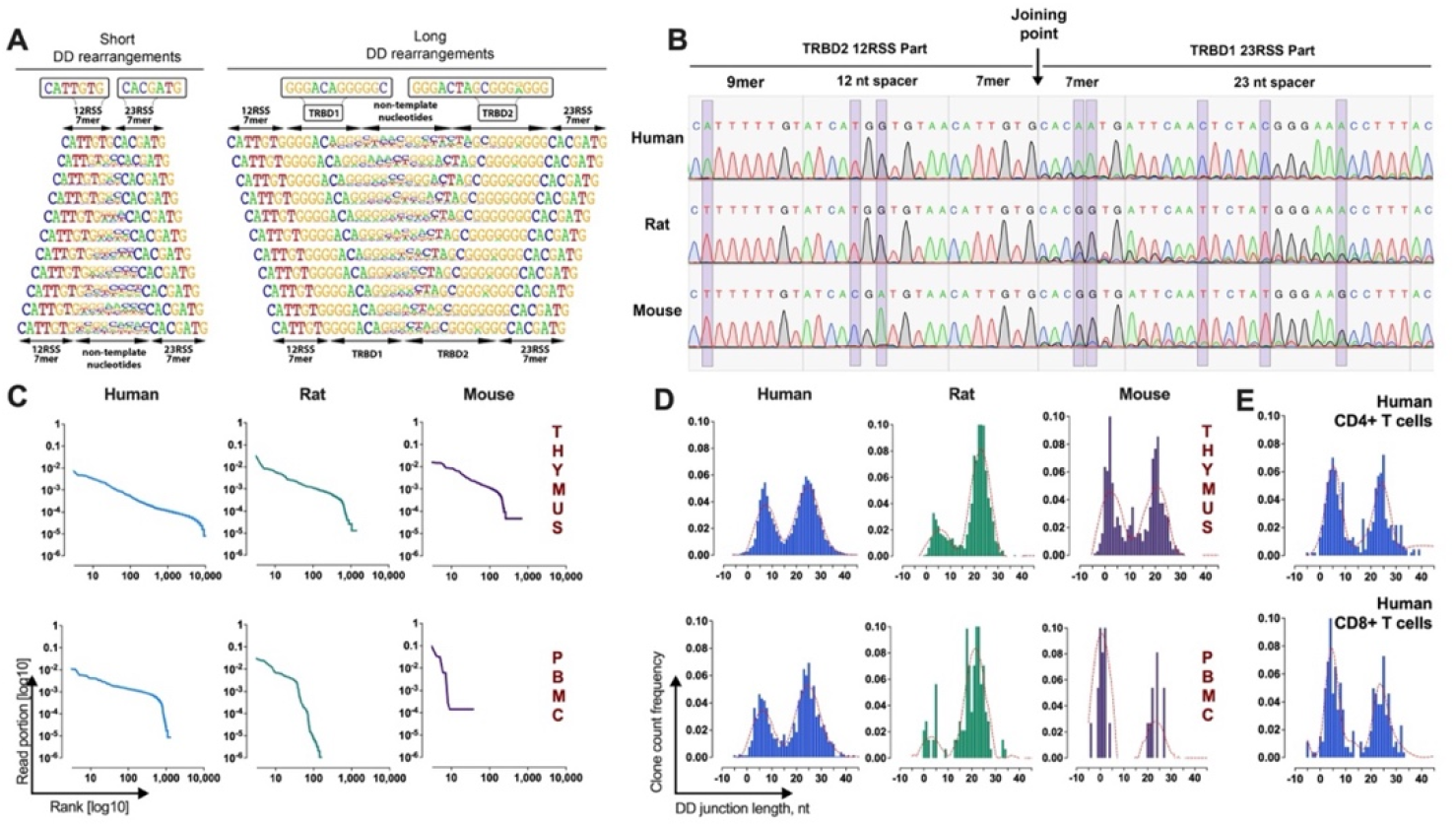
Identification and validation of TRB DD rearrangements in mammals ’lymphocytes. **A**. Frequency Logo diagram for short and long TRB DD rearrangements detected in human thymus DNA. **B**. Capillary sequencing of TRB D1 23RSS and D2 12RSS junction sequences from excision DNA circles from human, rat, and mouse thymes DNA. Violet bands show the position of variable nucleotides across the analyzed species. **C**. TRB DD rearrangements repertoire structures in human, rat, and mouse thymes and peripheral blood cells DNA. **D**. Bimodal distribution of TRB DD rearrangements junction length in human, rat, and mouse thymes and peripheral blood. **E**. Bimodal distribution of TRB DD rearrangements junction length in human helper and cytotoxic T cells.

Additional independent proof that TRB DD rearrangements produced by VDJ recombination machinery is the existence of its specific byproduct – T-cell excision circle DNA containing the junction of recombination signal sequences (RSS) from D1 and D2. Using PCR with primers specific for 23RSS adjacent to TRB D1 and 12RSS adjacent to TRB D2, we successfully detected TRB D1-D2 RSS junction in DNA from thymes of all analyzed species and then confirmed it by capillary sequencing (Figure 2B). All three sequences had expected structures containing 12RSS of the TRB D2 segment directly connected to 23RSS of the TRB D1 segment.

TRB DD repertoire profiling (Figure 2C) shows that DD rearrangements have a clonal distribution pattern similar to other TCR rearrangements being more abrupt for peripheral blood cells and smoother for the thymus. The obtained profile indicates that TRB DD rearrangements are present in normal cells and change their frequency following expansions and contractions of the cells that bear them. Genomic location, cell type specificity, nucleotide structure, RSS excision circles, and clonal distribution features of detected rearrangements prove that DD **rearrangements are indeed the product of the VDJ recombination process**.

### Bimodal distribution of TRB DD junction length

A deeper analysis of junction structures of TRB DD rearrangements shows that its length has bimodal distribution (Figure 2D). Thus, the DD rearrangements form two groups of “short” and “long” ones. The observed bimodality was characteristic for all analyzed species and for both thymes-derived and PBMC-derived DNA. The same distribution patterns have been observed in human helper T cells and cytotoxic T cells (Figure 2E). It means that both “short” and “long” DD rearrangements have been produced, most probably in the thymus before CD4/CD8 lineage commitment, i.e., at the stage of TCR gene formation, so some of them might contribute to TRB maturation. But being widely present in mature T cells, a large part of TRB DD rearrangements apparently just freeze in their incomplete state occupying the second, nonfunctional allele of TRB like another type of incomplete rearrangements -TRB DJ.

Next, we analyzed an additional group of DNA samples extracted from the peripheral blood of 15 healthy human individuals to measure the uniqueness of TRB DD rearrangements across the human population and the variability of TRB DD bimodal distribution (Figure 3).

**Figure 3.**
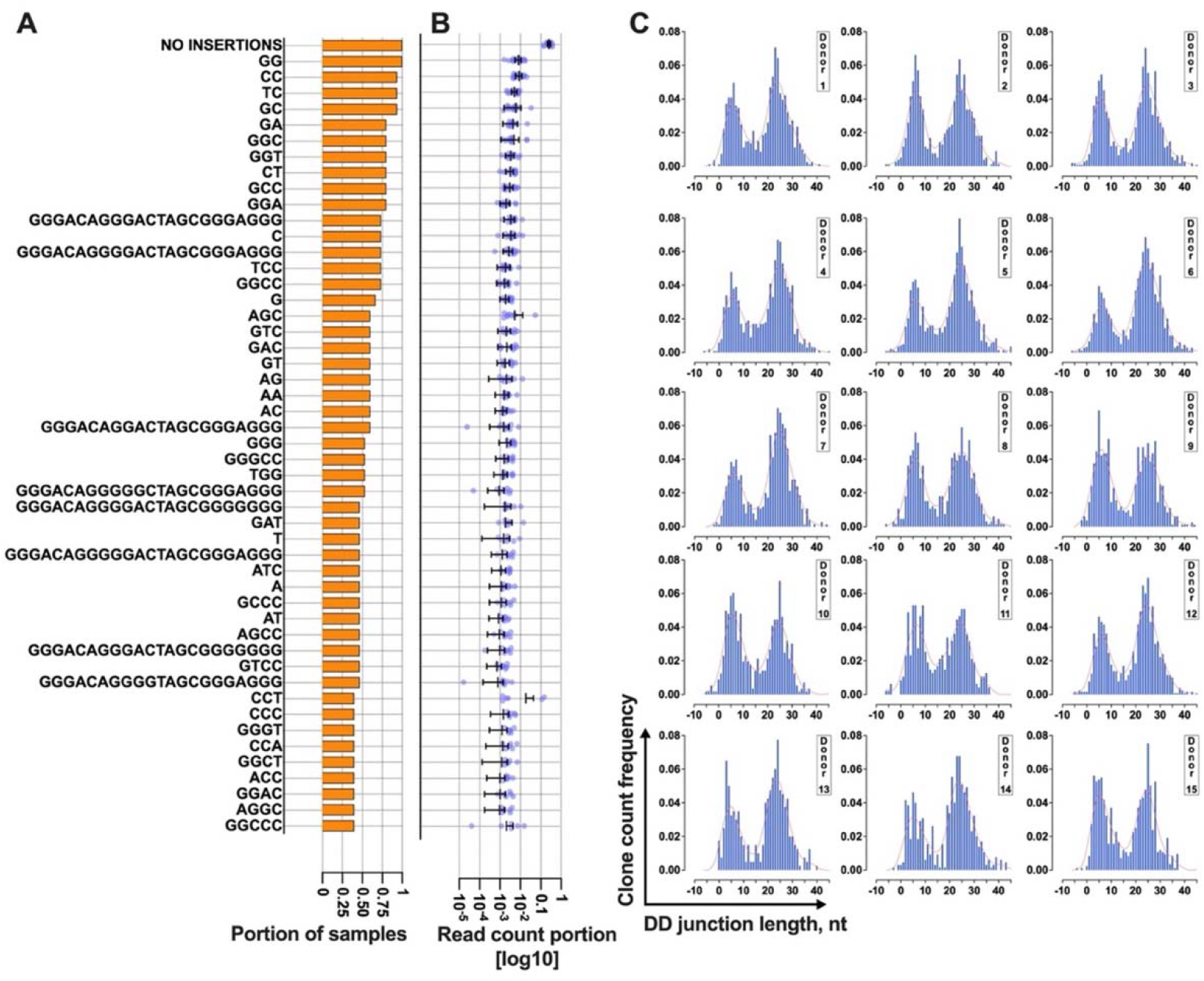
Bimodal distribution of TRB DD rearrangements across healthy individuals. **A**. The rate of cooccurrence of TRB DD rearrangements in DNA from different healthy donors. **B**. Read count frequencies of top 50 public DD clonotypes in T cells from 15 healthy donors. Bar and whiskers represent the mean with a 95% confidence interval. **C**. TRB DD rearrangement length distribution in T cells from 15 healthy donors.

The obtained results show that 16.6% of TRB DD rearrangements are public clonotypes present in two or more different samples (Figure 3A). It means they have a high generation probability and can be reproduced independently in different individuals. The most common rearrangements are those that lack inserted non-template nucleotides or have only 1-3 of them. This point is valid for both “short” and “long” DD rearrangements. The “short” rearrangement with zero added nucleotides is also the most frequent in each individual occupying one-quarter of all DD rearrangement-bearing cells (Figure 3B). Taken together, it indicates that this particular rearrangement is not random and is produced multiple times in each individual.

At the same time, the ratio of the number of “short” and “long” TRB DD can noticeably vary in the blood of different individuals (Figure 3C). Still, anyway, all have the same bimodal distribution pattern.

### Bimodality relates to DD recombination in two RSS sites

The DD junction lengths in two observed peaks had a substantial difference in ∼20 nucleotides. Meanwhile, there was no difference in the number of inserted non-template nucleotides (p=0.114, two-tailed Mann-Whitney test) but a significant difference in the number of missing nucleotides (p<0.0001, two-tailed Mann-Whitney test) of D1 and D2 segments. Moreover, the frequency of nucleotides matched to D1 and D2 in shorter rearrangements is not distinguishable from randomly generated insertions (Figure 2A). In other words, the short TRB DD junctions lack D1 and D2 fragments and are formed entirely by non-template nucleotide insertions. Thus, the main difference in DD junction length is based on the length of missing D segment fragments.

To understand if this bimodal distribution is unique across the VDJ rearrangements, we similarly analyzed TRB D1J1, and D2J2 rearrangement junctions, which do not contain DD junctions. Also, we analyzed previously known examples of DD rearrangements – D2-D3 rearrangements in the TRD locus. The distribution of DJ junction length was single modal (Figure 4C) in contrast to TRD DD, which shows the bimodal distribution like TRB DD (Figure 4B). Thus, the observed bi-modal distribution is most likely characteristic of DD rearrangements.

**Figure 4.**
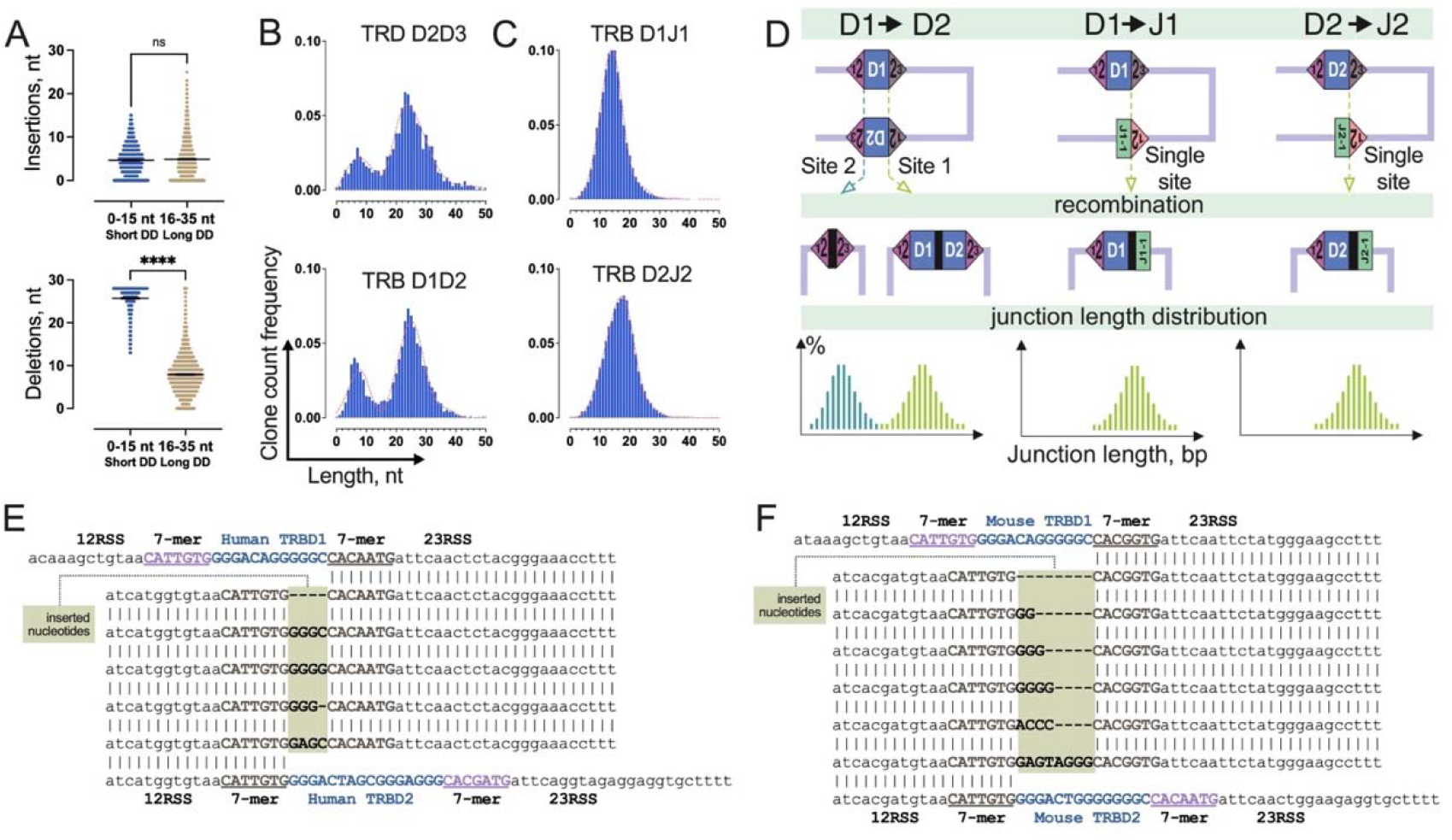
Two-site TRB DD recombination as a source of bimodality in DD junction length distribution. **A**. Comparison of the number of inserted non-template and deleted nucleotides in groups of “short” and “long” TRB DD rearrangements. Bar and whiskers represent the mean with a 95% confidence interval. **B**. Comparison of length distribution patterns of TRB DD and TRD DD rearrangements. Pooled 15 samples are displayed. **C**. Length distribution of TRB DJ rearrangements. Pooled 15 samples are shown. **D**. RSS pairs ’location differences during recombination between D1 and D2 segments and D and J segments in TRB locus. **E**. Inserted nucleotides in human TRB DD RSS junctions from excision circle DNA. **F**. Inserted nucleotides in mouse TRB DD RSS junctions from excision circle DNA.

The main structural similarity of D-D pairs of TRB and TRD loci is the presence of two putative sites of recombination (two 12/23 RSS combinations) in contrast to TRB D-J pairs which only have a single site (one 12/23 RSS combination) (Figure 4D). Therefore, the probable reason for bimodal distribution is the formation of two different DD RSS synapses with corresponding DNA cutting in two possible positions with the inclusion or exclusion of D1 and D2 segment sequences in the DD rearrangement structure. However, the orientation of RSSs in the second 12/23 RSS combinations site supposes that rearrangements retained in the genome should look rather like intrachromosomal RSS junctions without added nucleotides than normal diverse rearrangements. Indeed, the most frequent short TRB DD rearrangement (∼25%) is the one without any non-template nucleotides (Figure 3B), which have also been previously identified in mice (Curry and Schlissel, 2008; Hempel et al., 1998) due its high frequency.

To understand if there is a possibility of added nucleotides existing in the middle of RSS junctions, we additionally sequenced the amplicon of the human RSS junction (Figure 2B) on the Illumina machine. In obtained sequences, we indeed detected other non-canonical RSS junction structures presented in sequencing data 10 times less compared to previously detected classical RSS junction. These secondary RSS junctions appeared to have unexpected structures (Figure 4E). The difference from the classical ones was the presence of several additional nucleotides between RSSs, which is only partially matching to distal nucleotides of D1 and D2 segments. Similar secondary RSS junctions with additional nucleotide insertions were detected in a mouse (Figure 4D). Meanwhile, the sequences with full D1 and D2 fragments have not been identified among this data. Taken together, this indicates that the formation of “short” DD rearrangement occurs basically with the exact compliance to the 12/23 rule but with a kind of reversing of the biological sense and function of the target product (RSS-free rearrangement) and the byproduct of VDJ recombination (RSSs’ junction).

Thus, it can be concluded, that formation of observed non-canonical intrachromosomal RSS junctions with unusual random insertions is the mechanism of short DD rearrangements generation in contrast to long DD rearrangements which are producing in accordance with the classical VDJ recombination mechanism.

### The DD rearrangements are not a dead end of VDJ recombination

Analyzing the structure of RSSs remaining after recombination which are flanking DD rearrangements, we detected about 6% of corrupt RSS sites. They lacked one or more nucleotides at the ends adjacent to the DD junction, which is most probably the result of exonuclease activity during VDJ recombination. All defected RSSs were detected among “short” DD rearrangements, meaning this small fraction of it cannot be rearranged further. However, the rest 94% of DD rearrangements (both “short” and “long”) carry fully intact RSSs, so they can theoretically recombine with J segments using D2 23RSS and then with V segments using D1 12RSS.

To investigate this possibility, we performed the identification of TRB DD junctions firstly in incomplete DJ rearrangements (Figure 5A) and then in complete functional VDJ rearrangements (Figure 5B) from bulk PBMC of 15 healthy donors. TRB DJ was analyzed at the DNA level, and TRB VDJ was analyzed at the RNA level. The results show the unambiguous simultaneous presence of both D1 and D2 fragments in DJ and VDJ rearrangements. The most conservative estimation (7-mers of D1 and D2) of D1D2J2 in total D1J2 fraction was 17.7%, and of VD1D2J2 in functional VDJ2 fraction was 0.15% for bulk T cells.

**Figure 5.**
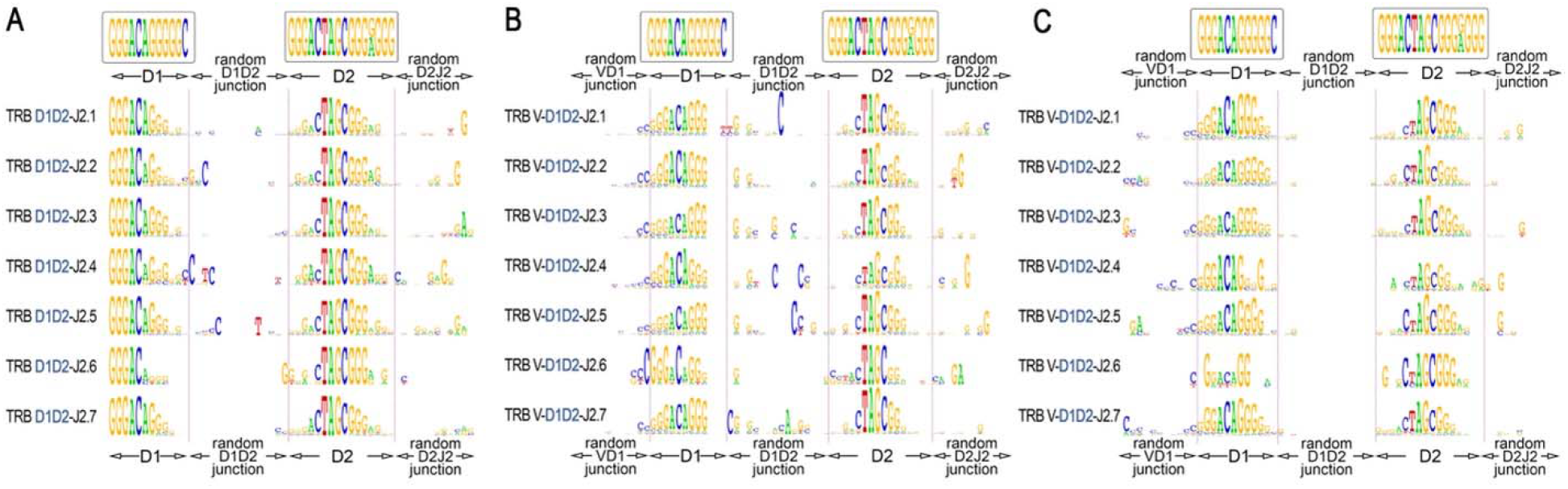
*TRB D1D2 junctions in the D1J2 and VDJ2 rearrangement structures*. ***A***. *Logo diagram of TRB D1D2J2 junctions among bulk PBMC (15 donors, data have been generated in this study)*. ***B***. *Logo diagram of TRB VD1D2J2 junctions among bulk PBMC (11 donors from previous research* (Sycheva et al., 2022), *data: PRJNA847436)*. ***C***. *Logo diagram of TRB VD1D2J2 junctions among memory (CD45RO) T cells. (11 donors from previous research* (Sycheva et al., 2022), *data: PRJNA847436). The results are displayed for pooled samples*.

Finally, we performed the same analysis for memory (CD45RO) T cells to understand if VDDJ rearrangement-derived TCR are genuinely functional and thus are capable of recognizing epitopes like conventional TCR. As a result, we successfully identified DD junctions in the structures of complete functional TCR VDJ rearrangements in memory T cells of 15 individuals (Figure 5C) at the level of 0.11% of the total functional VDJ2 fraction. This result shows that TCR genes containing hybrid DD segment have a functional impact on adaptive immunity having a capacity for antigens recognition.

Two orders of magnitude difference between DD junction recognition level in DJ and VDJ indicate that a major part of “long” DD junctions after the last stage of VDJ recombination cannot be detected directly. At the same time, the group of “short” DD junctions cannot be directly detected in VDJ rearrangements in principle since they lack D segment fragments. Thus, the real impact of DD rearrangements on the functional TCR generation might be dozens of times higher than we are able to detect for now.

## Discussion

The formation of adaptive immunity includes immune receptor diversity generation via VDJ recombination and subsequent negative and positive clonal selection waves in the thymus and second lymphoid organs. In this study, we analyzed noncanonical stage of VDJ recombination of the TRB locus and confirmed the existence of this process in humans for the first time. Our findings suggest that DD rearrangements in TRB locus are not marginal but rather regular part of TRB formation and diversification process in mammals.

We received the evidence that using two different recombination sites, this stage produces two groups of incomplete DD rearrangements: “short” and “long” ones. The first ones lack D segment fragments and contain only non-templated random nucleotides. The second one contains both D1 and D2 segment fragments and non-templated nucleotides. Most of the DD rearrangements from both groups retain intact external RSSs, which enable them to rearrange further as a hybrid D segment forming incomplete D1D2J2 and then complete VD1D2J2 rearrangements contributing to TRB diversity.

One additional step of diversification during DD joining should make derived mature TRB genes more heterogenic compared to classical ones. Furthermore, deleting J1/C1 segments cluster during DD recombination mechanically increases the frequency of J2 segments in TRB generation. This new knowledge could be the basis for the further improvement of the well-established TRB generation models (Dupic et al., 2019; Marcou et al., 2018) which was successfully applied for the computational TCR specificity evaluation (Pogorelyy et al., 2019, 2018).

The “short” DD group cannot be traced directly in complete TRB since they do not have recognizable parts of D segments. From this stage on, we can only hypothesize that they may contribute to shorter TRB fractions which are reported to participate in the immune response to common infections being abundant and public (de Greef and de Boer, 2021; Hou et al., 2019). The traces of the “long” group can be partially detected in incomplete and complete TRB rearrangements. Meanwhile, the “long” fraction is clearly contributing to antigen recognition being directly detected in memory T cell subsets in this study.

The presence of DD rearrangement in human, rat, and mouse genomes proves this stage of VDJ recombination to be conservative despite the slight differences in RSS signals in these species. DD rearrangements are also present in CD4 and CD8 T cells contributing to both arms of T cell immunity. This widespread distribution indicates an important role of this process in the development of adaptive immunity. Our results show that the TRB generation process is more complicated than previously thought, and thus the classical concept must be re-examined and corrected. It is obvious for now that there are at least two different paths for VDJ recombination, including the classical two-steps VDJ and alternative three-steps VDJ with DD rearrangement as a first step. The first one is probably quicker than the second one due to the number of required consecutive rearrangements.

In summary, this study analyzed almost unnoticed component of the TCR diversity generation process. For now, we do not completely understand how it affects T cell immunity in health and disease. However, we believe this new type of rearrangements is significant for both basic knowledge and applied research as a novel marker for clonality analysis (Semchenkova et al., 2022; Tirtakusuma et al., 2022) and minimal residual disease monitoring (Brüggemann et al., 2019; Nazarov et al., 2016) in lymphoid malignancies.

## Methods

### Sample collection and DNA isolation

The research was conducted according to the Declaration of Helsinki. All human subjects gave standard informed consent. The study was approved by the local ethical committee at Pirogov Russian National Research Medical University. In this study, we used DNA extracted from peripheral blood mononuclear cells (PBMC) from 15 healthy human individuals with the age range of 30-50 years, one wild-type mouse (Mus musculus) and one wild-type rat (Rattus norvegicus); DNA from the thymus of one human donor (16th donor, dissection of the thymus was a standard part of the heart surgery), DNA from thymes of the same mouse, and the same rat (Figure 1A); DNA from bulk CD8+ and bulk CD4+ human T cells. DNA from the RMS cell line and CD19+ human B cell fraction were used as a negative control.

### Isolation of PBMC and lymphocyte fraction

Peripheral blood mononuclear cells were isolated by Ficoll-Paque (Paneco, Russia) density gradient centrifugation according to a standard protocol. Bulk CD4+ and CD8+ T cells and CD19+ B cells were isolated from PBMC suspension using a magnetic separation approach with Dynabeads Positive Isolation Kits (Invitrogen, USA). DNA from all collected samples was isolated by FlexiGene DNA kit (Qiagen, Germany).

### TRB D1-D2 and DJ library preparation and sequencing

The library for high-throughput sequencing was obtained in two subsequent PCR reactions (Komkov et al., 2020). The first (target) 25 μl PCR contained 100 ng DNA isolated from PBMC, thymus or T cell subsets, 1X Turbo buffer, five units of HS Taq polymerase, 200 μM of each dNTP (all Evrogen, Russia), and 0.2 μM of each primer specific to genomic flanks of TRB D1-D2 junction for DD library (Figure 1B), primers flanking D and J segments for DJ library and primers flanking D2 and D3 for TRD D2-D3 library (all MiLaboratories, USA). The amplification profile was 94°C for 3 min, followed by 10 cycles of 94°C for 20 s, 57°C for 90 s, and 72°C for 40 s, followed additional 15 cycles of 94°C for 20 s and 72°C for 90 s. All PCR cycles were performed with Ramp 0.5°C/s. Six replicates were obtained for each PBMC and thymus sample to reach the average sample size equal to ∼100,000 analyzing cells (600 ng input DNA total per sample).

Obtained amplicons were purified using 1V AmPure XP beads (Beckman Coulter) and used as a template for the second PCR. The second (indexing) 25 μl PCR contained 1X Turbo buffer, 2.5 units of HS Taq polymerase, 200 μM of each dNTP, and 1 μl of Unique dual indexes primers (Illumina, USA). The amplification profile was as follows: 94°C for 3 min, followed by 15 cycles of 94°C for 20 s, 57°C for 20 s, and 72°C for 40 s. Obtained amplicons were purified using 0.8V AmPure XP beads (Beckman Coulter, USA), pooled, and sequenced on Illumina NextSeq or MiSeq machine with coverage ∼100,000 reads per sample (Figure 1C).

### TRB D1-D2 RSS signal junction identification

RSS signal junction detection was performed using PCR with subsequent capillary sequencing. To obtain the target PCR product, 150 ng of DNA isolated from the thymus (human, rat, and mouse) was used as a template in a 25 μl PCR reaction containing 1X Turbo buffer, 2.5 units of HS Taq polymerase, 200 μM of each dNTP and 0.2 μM of each primer specific to RSS junction flanks (Figure 1B, Figure 2B). The amplification profile was 94°C for 3 min, followed by 45 cycles of 20 s of 94°C, 20 s of 58°C and 40 s of 72°C. RSS signal junctions for each analyzed species were amplified in separate PCR. Capillary sequences of the target PCR products were obtained as a service (Evrogen, Russia).

### TRB D1-D2 and DJ repertoire reconstruction and analysis

TRB DD, DJ, and TRD DD rearrangements were extracted from fastq files using MiXCR3.0 software (Bolotin et al., 2015) with additional genomic reference, containing TRD D2, D3, TRB D1, D2, and J segments with their genomic flanks from IMGT database (Giudicelli et al., 2005). VDJtools software (Shugay et al., 2015) was used for subsequent post-analysis: clonotype diversity calculation, clonotype uniqueness analysis, and detection of inserted and deleted nucleotides in the DD junction. Statistical analysis and plotting were performed using GraphPad Prism 9.0. To detect DDJ rearrangements, the standard terminal “grep” function was used to search for 7 nt k-mers (ggactag, gactagc, actagcg, ctagcgg, tagcggg, agcgggg, agcggga) specific for D2 segments in D1-J2 junction fragment. The sequences containing target k-mers were subjected to multiple alignment using SnapGene software with MAFFT v7.471 algorithm (Katoh and Standley, 2013). The LOGO diagrams were produced by WebLogo software (Crooks et al., 2004) using aligned sequences for DDJ and non-aligned ones for DD rearrangements.

### VDDJ rearrangements detection

TRB VDJ rearrangements were extracted from previously published deep TCR-seq dataset (Sycheva et al., 2022) for bulk PBMCs of 11 healthy donors (24-60 years old) before vaccination (PRJNA847436) using MiXCR 3.0 with default parameters. Obtained clonotype tables were converted into VDJtools format using “Convert” function with “-S mixcr” parameter. The level of quantitative bias was checked using iROAR software (Smirnova et al., 2023). The TRBJ2 containing clonotypes with detected D1 segment were extracted from the analyzed dataset using “FilterBySegment” function and then were separated to functional and nonfunctional clonotype tables using the function “FilterNonFunctional”. The final detection, aligning, and displaying of VDDJ2 rearrangements were performed as described above for DDJ2 rearrangements.

## Ethic declarations

The research was conducted according to the Declaration of Helsinki. All human subjects gave standard informed consent. The study was approved by the local ethical committee.

## Competing interests

The sequences of primers specific for DD and DJ rearrangements used in this study are intellectual property of MiLaboratories LLC., USA. Authors declare no other competing interest.

## Data availability

The sequencing data generated in this study has been deposited to SRA NCBI under accession number PRJNA952099. Previously published TCR-seq data19 used in this study are available under accession number PRJNA847436.

## Authors’ contributions

AYK – the idea, study design, funding, data analysis, manuscript drafting; AOS, AMM, LDB and IVK – performed the experiments and data analysis; YBL, IZM, DMC – resources, samples, mentorship, manuscript drafting. All authors reviewed and approved the manuscript.

## Notes

### Competing Interest Statement

The authors have declared no competing interest.

### Summary of Updates

Title and abstract are edited

